# Cell-lineage selection interacts with development and life history to shape the rates and identity of heritable mutations

**DOI:** 10.64898/2026.11.02.763612

**Authors:** Paco Majic, Malvika Srivastava, Justin Crocker

## Abstract

Mutation rates determine the supply of heritable variation and are central to evolutionary theory, yet they are usually treated as properties of molecular mutational processes. We hypothesised that mutation rates are also shaped by selection among cell lineages within organisms and that the strength of this process depends on developmental and life-history traits. To test the extent of this idea, we combined mathematical modelling with experiments in *Drosophila melanogaster* and treated mutation rates as emergent properties of interactions among molecular mutagenesis, development, life history, and cell-lineage selection. The model predicts that even modest effects of mutations on cellular fitness can substantially alter the probability that mutations are transmitted, and that these effects depend strongly on developmental architecture, germline dynamics, and time to reproduction. In *Drosophila*, we experimentally demonstrate biased transmission of a mutation affecting cellular proliferation and show that this bias changes over the reproductive lifespan of the individual. Extending the model genome-wide reveals that the cellular distribution of fitness effects and its interaction with developmental parameters influence both the estimated mutation rate and the composition of transmitted mutations, producing transmission biases that favour mutations with particular cellular fitness effects. These results suggest that developmental and life-history parameters determine how cellular fitness effects translate into transmission bias; therefore, estimates of heritable mutational input can vary from species to species even under constant molecular mutagenesis. By filtering mutations before they enter populations, developmental and life-history evolution can modify both the rate and the nature of heritable variation and thereby influence evolutionary trajectories.

**Significance Statement:** Mutations fuel evolution, and their rate can influence the pace of species diversification. While the mutation rates of populations are influenced by molecular rates of DNA changes, the context in which such changes first arise also determines how many and which mutations enter a population’s gene pool. In multicellular species, such context is the body of an individual organism. Thus, mutation rate differences between populations may result from differences in developmental traits, and also by how mutations affect the fitness of cells within their developmental environments. This highlights how, in evolution, organisms are not simply the object of selection, they also help determine the pace and direction of genotypic evolution by influencing how many and which mutations are passed on.

## Introduction

Mutations provide the raw material for heritable variation in populations, and their rates can define the tempo and direction of adaptation. But before entering the gene pool of a population, all mutations first arise in a single cell in an organismal context. In the case of multicellular organisms, this context is the microenvironment constructed by the populations of cells that make up the organism. Because mutations may affect how cells thrive and proliferate within those microenvironments, i.e. their fitness, the first selective sieve for any mutation is the organism itself.

Mutations that increase cell fitness eventually become overrepresented within tissues, while those that decrease it become underrepresented. At the somatic level, competition between cell lineages is often studied as a mechanism of tissue-level quality control (1-3), it has been especially prominent particularly in the context of cancer and senecense, both for medical reasons (4-8) and within evolutionary frameworks (9-11), while it has also been proposed to positively affect individual fitness (12-14) with the potential of steering evolutionary trajectories (15, 16). When cell-lineage selection occurs within the germline, its consequences extend beyond the organism itself: by biasing which mutations are represented in gametes, germline cell-lineage selection can systematically shape the allelic input into populations (17-21). Empirical evidence for how cell-level selection may lead to such bias exists across diverse taxa and developmental contexts. Experimental support comes from *Drosophila*, both in ovaries (22) and testis (23), while additional evidence includes cases in which inheritance is affected by selection in plants (24), and mitochondrial mutations in mammals (25, 26). In humans, some of the best-studied cases of cell-lineage selection in the male germline involve conditions such as Apert and Noonan syndromes (27-29), while very recent studies using high-resolution single-cell sequencing of exomes from human sperm samples have shown that positive selection in the male germline drives clonal expansions across a broad spectrum of genes (30, 31).

The rate at which new mutations can enter the gene pool of a population (*µ*_*pop*_) is the result of an interaction between the rate at which new mutations arise at the molecular level (*µ*_*mol*_) and the life-history traits of an individual (32-34). Because germline cell-lineage selection biases which mutations ultimately enter the gene pool, it can influence *µ*_*pop*_ and, consequently, the amount of genetic variation generated each generation. Yet, current explanations for mutation-rate variation at the level of species or populations seldom, if ever, consider the context of cell-lineage selection. These explanations tend to largely treat mutation rates as properties of molecular mutational processes, modulated by organismal traits such as generation time and numbers of cell divisions (as reviewed in (33, 35-39)). This perspective underlies explanations for major patterns in molecular evolution, including the drift-barrier hypothesis (37, 40-42), paternal biases (43-45), and the generation-time effect (46, 47). But as was also noted by Choi et al (27) for mutations causing Apert syndrome, elevated mutation incidence is likely driven not by increased molecular mutation rates, but by selective advantages conferred during spermatogenesis. Indeed, certain mutational patterns that can be attributed to variation in mutagenesis, may actually result from cell-lineage selection more broadly across the genome (21). Extending this principle farther, a selective bias at the cellular level across loci can bias how the causes of mutation rate variation between and within species are interpreted.

The selective consequences for mutations within organisms are shaped by how they affect intrinsic cellular processes (48), but also by traits of those organism within which they arise. Cells are exposed to pressures from their tissue-specific microenvironments which may impose survival and proliferative challenges as well as competition for nutrients, signaling molecules or physical space (49, 50). These selective dynamics may be further influenced by cellular demographic parameters such as tissue size, the amount of genetic variation among cells, and their spatial organization (51-53). In addition, the timing and duration of developmental events may modify both the intensity and the targets of selection (54) thus altering which cellular pathways and processes that are challenged at different stages of development. Consequently, we expect that as species evolve diverse morphologies and life-history strategies, the selective conditions experienced by cell lineages within organisms also change, potentially increasing or decreasing the importance of cell-lineage selection within the germline. Thus, while cell-lineage selection may be a major contribution to the patterns of genetic diversification of certain developmental modes, it may be irrelevant for others. We therefore hypothesized that the influence of cell-lineage selection on both the number and identity of mutations that ultimately enter populations depends on organismal biology, including developmental architecture and life-history traits.

To begin exploring this hypothesis, we combined mathematical modelling with experiments in *Drosophila melanogaster*. Building on the theoretical framework of Otto and Orive (18), which examined the consequences of intraorganismal selection for mutation load, we extend this perspective by treating mutation rates themselves as emergent properties arising from interactions among cell-lineage selection, development, and life history. Our results highlight the importance of these interactions and they point to the need of understanding species-specific differences in the cellular distribution of fitness effects, a parameter that remains largely unexplored empirically, particularly in multicellular organisms and in comparative studies. More conceptually, we propose that organisms, by imposing the first selective filter on new mutations, play an active role in shaping the heritable variation available for evolutionary change within their own lineages.

## Results

### Cellular fitness effects interact with development and life-history to shape per-locus mutation rates

We extended our previous developmental model of mutation-rate evolution (32), which distinguishes the molecular mutation rate (*µ*_*mol*_) from the generational mutation rate (*µ*_*pop*_), defined as the probability that a mutation arising during development is represented among the gametes produced by an individual (Fig. 1A; Methods). In the original model, *µ*_*pop*_ depends not only on *µ*_*mol*_, but also on developmental parameters including the rate of cell division (*C*), developmental duration (*τ*) and the timing of germline sequestration (*γ*). Here, following previous theoretical work (18), we incorporated cell-lineage selection by assigning each mutation a cellular fitness effect (*β*), allowing mutations to alter the proliferative success of the lineage in which they arise (Fig. 1A). We further considered the extension of the model that includes a maturation phase between development and reproduction, during which a constant pool of germline stem cells undergoes turnover (32). Turnover occurs at rate *d* over a maturation period of duration *l*, while mutations influence stem-cell competition through additional fitness parameters affecting the rate at which mutant germline stem cells are lost, *β*_*d*_, or replenished, *β*_*l*_ (Fig. 1A; Methods). For the developmental phase, some versions of our model considered *µ*_*mol*_ to either reflect mutations per locus per cell divisions, capturing the rate of mutations of replicative origin and which is the baseline we use throughout this manuscript, or unit time, capturing the clock-like accumulation of mutations caused by DNA damage and repair (55). For the maturation phase models, we only considered the latter. For simplicity, we also assumed that the fitness effects of mutations on a specific locus, did not interact with mutations at other loci.

**Figure 1.**
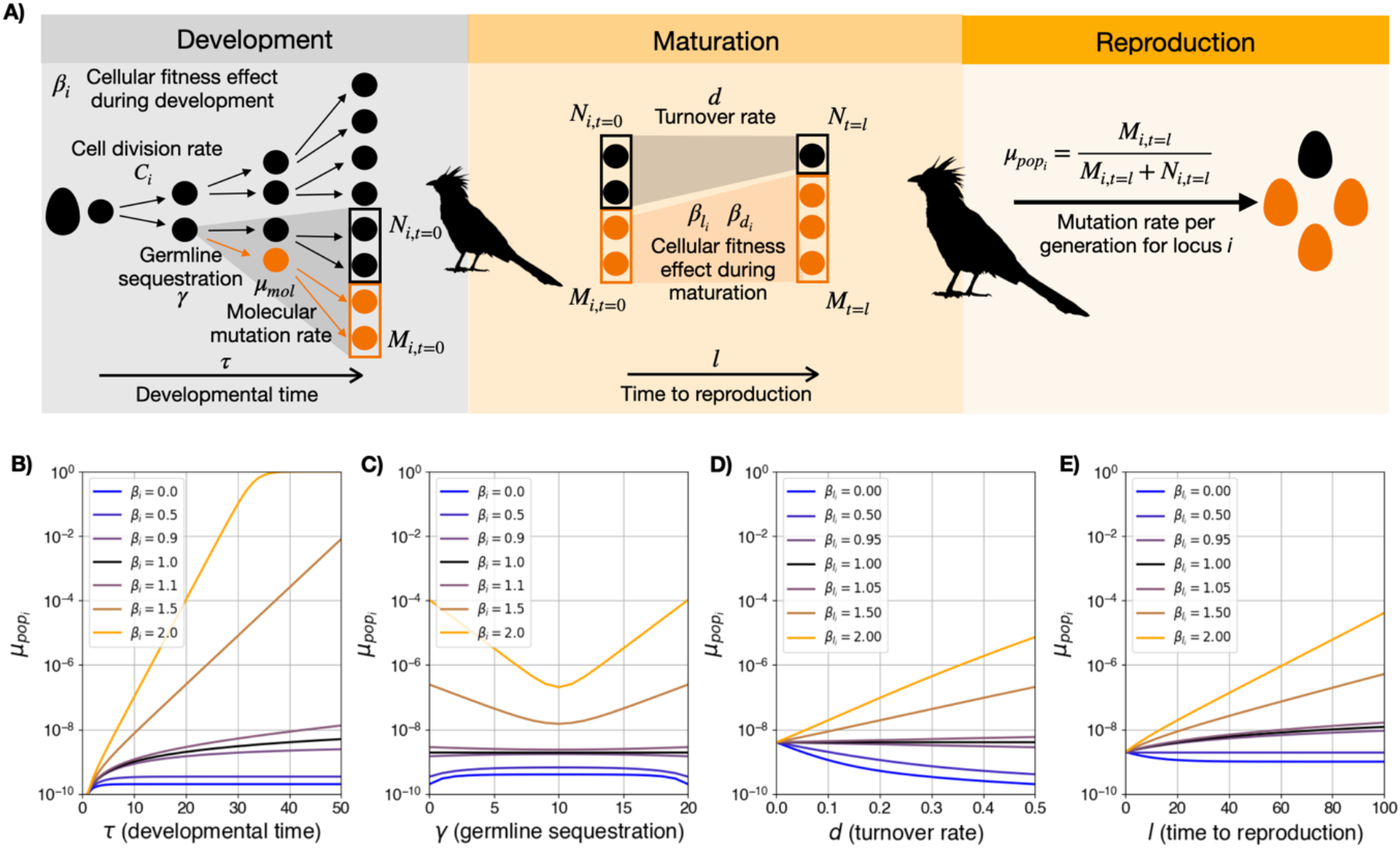
Developmental and life-history parameters interact with cell-lineage selection to shape generational mutation rates. **(A)**Schematic of the model. During development, cells divide at rate *C* for a duration *τ*. Germline sequestration occurs at cellular generation *γ*, at which a single cell founds the germline lineage. Mutations arise during cell division at rate *μ*_mol_ per locus and have a locus-specific cellular fitness effect *β*_i_. At the end of development, mutant and non-mutant cell numbers are *M*_i_ and *N*_i_, respectively. During the maturation phase, the germline stem-cell population may undergo turnover at rate *d*. Mutations arising during this phase occur per unit time and have fitness effect *β*_*l*_ or *β*_*d*_. At reproduction, the generational mutation rate for locus *i* 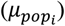 is calculated as the probability of sampling a mutant cell from the pool of gamete-producing cells. **(B)** 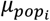 as a function of developmental duration (*τ*) for different values of *β*_i_. *C* was fixed at 1 and *μ*_mol_ was fixed at 1 × 10^−10^ mutations per locus per cell division, close to empirical estimates for mammalian germline cells (127). In this case *γ* and *l* were held as 0, meaning there was no influence from germline seqeustration or time to reproduction. **(C)** 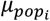 as a function of germline sequestration time (*γ*) for different values of *β*_i_. *τ* was held at 20 and all other parameters were the same as in B. **(D)** 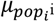 as a function of the rate of germline stem cell turnover (*d*) for different values of 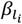!. Here, time to reproduction (*l*) was held constant at l = 10, *γ* was held at 0 and all other parameters were the same as in C. **(E)** 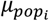 as a function of time to reproduction (*l*) for different values of 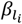. *d* was held at 0.1 and all other parameters were the same as in D. **(F)** 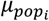 as a function of the fitness effect of a mutation on locus i before (*β*_*i*_) and after 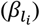 the end of development and time to reproduction *l*. All developmental parameters are here set to their baseline values (*β*_*i*_ = 1, *µ*_*mol*_ = 1*x*10^−10^, *C* = 1, *γ*=1, *τ* = 20), while stem cell turnover, *d*, was set to *d* = 0.1.

In our model, varying the proliferative effect *β* of a mutation on locus *i* (*β*_*i*_) can drastically change the effective *µ*_*pop*_ for that locus 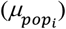 even when *µ*_*mol*_ is constant, such that higher values of *β*_*i*_ imply higher 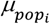 (Fig. 1B-E). Interestingly, even mild consequences for cellular proliferation may have major consequences for the rate at which new mutations enter the gene pool of populations. For example, in the case of *Cτ* = 20 (representing 2^20^ cells, roughly the number of ovarian follicles an infant is born with (56) or half of the number of germ cells in the ovary of cows by the third trimester of pregnancy (57)), a mere 1% increase (*β*_*i*_ = 1.01) or decrease (*β*_*i*_ = 0.99) in cell proliferative capacity may result in a ∼6% change in 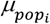 relative to a neutral case (*β*_*i*_ = 1). While more pronounced fitness effects of 5% increase or decrease (*β*_*i*_ = 1.05 and *β*_*i*_ = 0.95, respecitvely) could result in shifts in 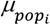 of ∼25%. Furthermore, higher values of *Cτ* - which would result in a greater number of cells, and thus larger organisms (58)- tend to produce larger differences in 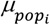 for large *β*_*I*_ (Fig. 1B). On the other hand, when *β*_*i*_ is small, *Cτ* does not have a strong effect on the mutation rate. This is true if we consider mutations arising during development to be of replicative origin and thus track cell divisions (Fig. 1B, Fig. S1A), but also if mutations are considered to result from DNA damage and repair, in which case mutations track units of time (Fig. S1B). The effect of *β*_*i*_ on 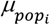 therefore depends on the developmental parameters *C* and *τ*.

Varying the timing of germline sequestration (*γ*) also altered the effect of cell-lineage selection on 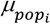. Across the explored parameter space, the interaction between *γ* and *β*_*i*_ was strongly non-linear, with selection having the weakest effect when germline sequestration occurred midway through development and the strongest effects when sequestration occurred either very early or very late (Fig. 1C). This pattern arises because *γ* determines how long differences in cellular proliferation can influence the relative representation of mutant and non-mutant cells that contribute to the germline. When sequestration occurs very early or very late, selection acting on cell proliferation has a greater opportunity to alter the composition of the lineage that ultimately contributes to the germline, whereas sequestration at an intermediate developmental stage reduces this effect. We also allowed mutations to have distinct fitness effects before and after germline sequestration by differentiating between a zygotic and germline fitness effect of a same mutation on locus *i* (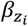 and 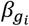 respectively, see Methods). This captured a context dependence of the fitness effect of one same mutation, which further expanded the range of possible dynamics in terms of how cell-lineage selection interacts with germline sequestration to shape 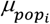 (Fig. S1C).

Lastly, time to maturation (*l*) can also alter the effect of cell-lineage selection on 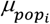. When considering the effect of cell-lineage selection in a maturation phase before reproduction, whenever there is a turnover of cells and mutations at locus *i* do affect the loss or replenishment rates of cells, by respectively affecting 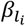 or 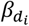, a positive effect of mutations on the fitness of cells increases the overall 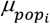 while deleterious mutations decrease it closer to *µ*_*mol*_ (Fig. 1D, Fig. S1D). Importantly, this effect is more pronounced for individuals that take longer to reproduce (Fig. 1E, Fig. S1E). In the case when there is no turnover of cells during the lifetime of an individual (*d* = 0) or when mutations are neutral 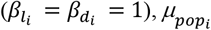 only changes due to the accumulation of new mutations (Fig. 1E, S1E).

Together, these results demonstrate that developmental architecture, germline organisation and post-developmental stem-cell dynamics interact with cell-lineage selection to determine the effective mutation rate of individual loci.

### Inference of the cellular fitness effect of genetic variants during *Drosophila* development

Although there is strong empirical evidence supporting the idea that selection at the cellular level in the germline can bias heritability of new mutations, a direct empirical validation that changes in cell proliferation may bias transmission, as in our model, is still lacking. To fill this gap, we conducted experiments on *Drosophila melanogaster*. Specifically, we considered alleles of two genes with different expected effects on cellular fitness. On the one hand, we considered the gene *trithorax-related (trr)*, which encodes for a methylase targetting lysine 4 of histone H3 and is therefore involved in chromatin function and patterns of gene regulation (59-61), Previous studies have shown that the mutational impairment of the function of this enzyme influences cell proliferation (62), leading to larger flies (63). On the other, we considered the classic *w*^1118^ allele of the gene *white*, originally described by Morgan (64), which encodes an ATP-binding cassette (ABC) transporter involved in pigment precursor transport typically used as a phenotypically “neutral” marker in *Drosophila* studies (but see (65)).

Our experiments, illustrated in figure 2A, consisted on generating mutant clones by inducing mitotic recombination through the FLP/FRT technique (66) on heterozygous F1 embryos that carried a sub-functional allele of *trr* (TrrCA,(63)) and a wild-type version of the gene (TrrWT) on a chromosome marked with a *red fluorescent protein, mRFP* (see Methods). As a result, within a single mosaic embryo, some cells are homozygotic for TrrCA, homozygotic for TrrWT, or heterozygotic with a copy of each allele. Because an impaired *trr* activity increases the rates of cell proliferation (62), we expected that the TrrCA allele would be overrepresented in the germline and thus be more prone to be inherited than the TrrWT allele. To test this, we produced single crosses of females during the development of which we had induced mitotic recombination with males that had a wild-type version of *trr* and no fluorescent marker. For each female we then measured the number of fluorescent F2 offspring. Consistent with our hypothesis, when mosaicism was induced early in development, a median of ∼56% of offspring carried the mutant allele (were not fluorescent), a fraction that was significantly higher than the cases when mosaicism was not induced on the control TrrCA/TrrWT heterozygotes (Fig. 2B). When we induced the FLP/FRT that had two copies of TrrWT, but was heterozygotic for *w*^1118^, which is not expected to affect cell proliferation in the germline (*β*_w1118/ w1118_ = *β*_+/w1118_ = *β*_*+/+*_), the *w*^1118^ mutation from the mother was inherited in similar proportions to the allele carrying the fluorescent marker (“TrrWT (HS)” in Fig. 2B).

**Figure 2.**
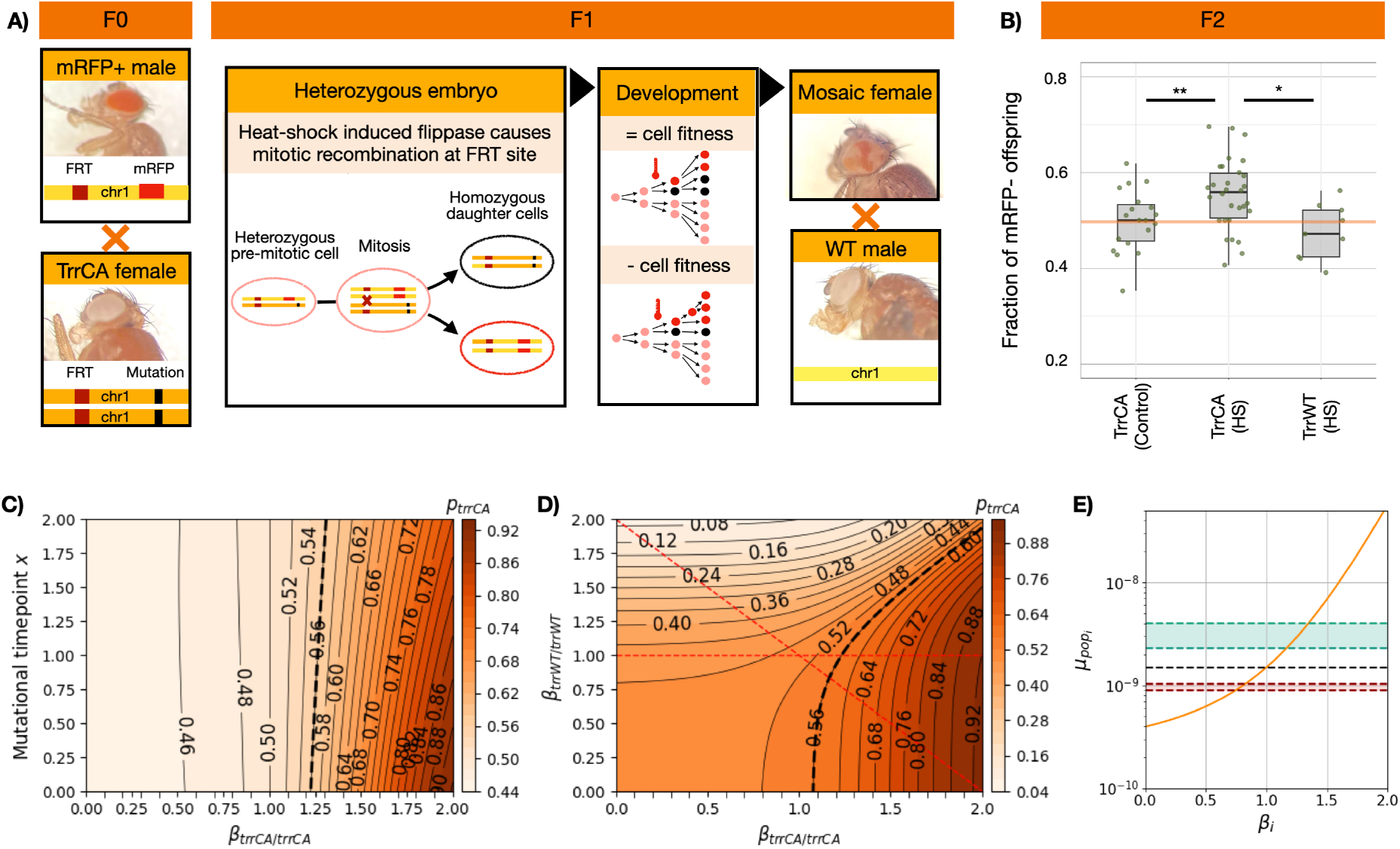
Cell-lineage selection in the germline of female *Drosophila* mosaics biases allelic inheritance. **(A)** Illustration of our FLP/FRT experiment. Mitotic clones were generated using a standard hsFLP/FRT system in *Drosophila*. Flies carrying mRFP, hsFLP, and FRT sites crossed to lines bearing also an FRT site associated to a *trr*^CA^ or *w*^1118^ mutation to produce heterozygous embryos. Embryos collected for 1 hour, were heat-shocked within 4 hours after laid at 37 °C for 1 hour to induce FLP-mediated recombination and clone formation. Mosaic and non-heat-shocked control virgin females were crossed individually to *w*^1118^ males. Offspring were scored for mRFP fluorescence to estimate transmission of the mRFP-positive versus recombinant (*trr*^CA^, *w*^1118^) allele. **(B)** Fraction of offspring from mosaic females that did not carry the marker mRFP allele from vials for a case in which FLP was induced by heat shock (HS) or not (Control) and whenever the alternative to mRFP was either a mutant TrrCA or TrrWT. **(C)** Probability of sampling the TrrCA allele upon reproduction (*p*_trrCA_) depending on the different possible values of the cellular fitness effect of that mutation in homozigosity, *β*_trrCA/trrCA,_ and the developmental timepoint at which flippase-iduced recombination happens (Methods). For this results *β*_trrWT/trrWT_ was assumed neutral relative to the backgrund heterozygous cells. The thick dashed black cline indicates the emprical expectation of *p*_trrCA_ given the results in (B). **(D)** *p*_trrCA_ depending on the different possible values of the cellular fitness effect of TrrCA and TrrWT in homozigosity, *β*_trrCA/trrCA and_ *β*_trrWT/trrWT_, respecitvely. In this case *x* = 0, but see figure S2 for the case in which *x* = 1 and *x* = 2. The thick dashed black cline indicates the emprical expectation of *p*_trrCA_ given the results in (B). The horizontal red dashed lines represents the case in which *β*_trrWT/trrWT_ is neutral relative to the backgrund heterozygous cells, as in C, while the diagonal red dashed line indicates the case when the cellular fitness effects of the TrrCA allele are additive. **(E)** Estimation of generational mutation rates, 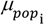, for a locus *i* at which mutations cause cellular fitness effects during development, *β*, similar as those we estimated for our *Drosphila* experiment. The black dashed line indicates the expected 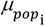 under neutrality (*β*_*i*_ = 1), while the green shaded area indicates the range for estimated beneficial fitness effects and the red shaded one indicates the range for deleterious effects (see Table S1). The parameters considered were *C*τ = 15 cycles of cell division, *μ*_mol_ was fixed at 1 × 10^−10^ mutations per locus per cell division and *γ* = 6, because although the formation of the population of nuclei that will develop into pole cells of the germline is after 9 rounds of nuclear divisions (123), the founding population of the germline is composed of roughly 10 nuclei, which would have undergone approximately 6 cycles of nuclear divisions since the zygote.

We adapted our model to our experimental setup to estimate what are empirically plausible values for the cellular fitness effect of cells carrying the TrrCA allele, *β*_trrCA/trrCA,_ based on the proportion of TrrCA mutant among the F2 offspring. Specifically, we incorporated what is known about germline development in female *Drosophil*a (Methods). In our experiment, mitotic recombination was induced early in development during the window of time in which primordial germ cells (PGCs) are expected to undergo two division cycles (Methods). Because the exact cell division cycle at which recombination happens, *x*, is expected to vary from embryo to embryo, we considered values of *x* to range from 0 to 2 cell divisions. Assuming that TrrWT is dominant in its fitness effects in heterozygotic cells (*β*_trrWT/trrCA_ = *β*_trrWT/trrWT_), we inferred *β*_trrCA/trrCA_ values that reproduced the experimentally measured effective transmission probability of the TrrCA allele (Fig. 2B). For a probability, *p*_trrCA_, of 0.56, *β*_trrCA/trrCA_ ∼1.2 (Fig. 2C, Table S1). Relaxing the assumption that TrrWT has a dominant fitness effect (*β*_trrWT/trrWT_ ≠ 1), altered the inferred values of *β*_trrCA/trrCA_ for *p*_trrCA_ = 0.56 (Fig. 2D, Fig. S2), such that for an additive scenario (red diagonal in Fig. 2D), *β*_trrCA/trrCA_ ranges from 1.16 to 1.21, and *β*_trrCA/trrCA_ from 0.79 to 0.84, depending on *x* (Table S1).

To explore the potential implications of these experimentally observed transmission biases and their associated cellular fitness effects for generational mutation rates, we then used our original developmental model (Fig. 1A) with developmental parameters similar to those of *Drosophila* (*Cτ* = 15, and *γ* = 6, see Methods). Values of *β* comparable to those of our estimates for *β*_trrCA/trrCA_ (1.16 < *β*_trrCA/trrCA *<*_ 1.31, Table S1) resulted in a 1.51- to 2.38-fold increase in 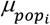 relative to a locus at which mutations are neutral at the cellular level (Fig. 2E, Table S1). In contrast, values of *β* comparable with the deleterious effects estimated for *β*_trrWT/trrWT_ result in a 0.65-to 0.70-fold change (Fig. 2E, Table S1).

We note that these inferred values of cellular fitness effects should be taken as approximations that, given the experimental design, primarily reflect the effective transmission probability of engineered alleles in a mosaic germline, rather than the fate of naturally arising de novo mutations.

### Variation of cellular fitness effects over lifetime

Interestingly, the transmission bias of the TrrCA allele reversed over the lifetime of mosaic females. When we followed the transmission of the TrrCA over the lifetime of individual females transferred weekly to new vials (Fig. 3A), we noted that although these females produced progeny biased toward the mutant allele immediately after pupation (Fig. 2B), the bias shifted in favor of the TrrWT marked with mRFP as those females aged (Fig. 3B). This reversal is consistent with a change of the cellular fitness effect of TrrCA over the lifespan of individual flies, such that while *β*_trrCA/trrCA_> 1 during development, 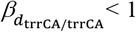 in the adult female germline. In contrast, as expected, TrrWT associated to *w*^1118^ retained a neutral cellular fitness effect throughout development and the lifespan of each female fly (Fig. 3C). We adapted our model to account for time to reproduction following development and applied it to these experiments (Methods). We estimated the expected proportion of TrrCA alleles in offspring for different values of *l* and different cellular fitness effect of the TrrCA in the adult female germline, 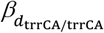 (Fig. 3D). Based on the effective transmission probability of TrrCA alleles in F2 offspring measured each week and assuming 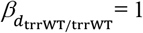, we inferred a 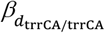 ranging from 0.76 to 0.84 (Fig. S3A).

**Figure 3.**
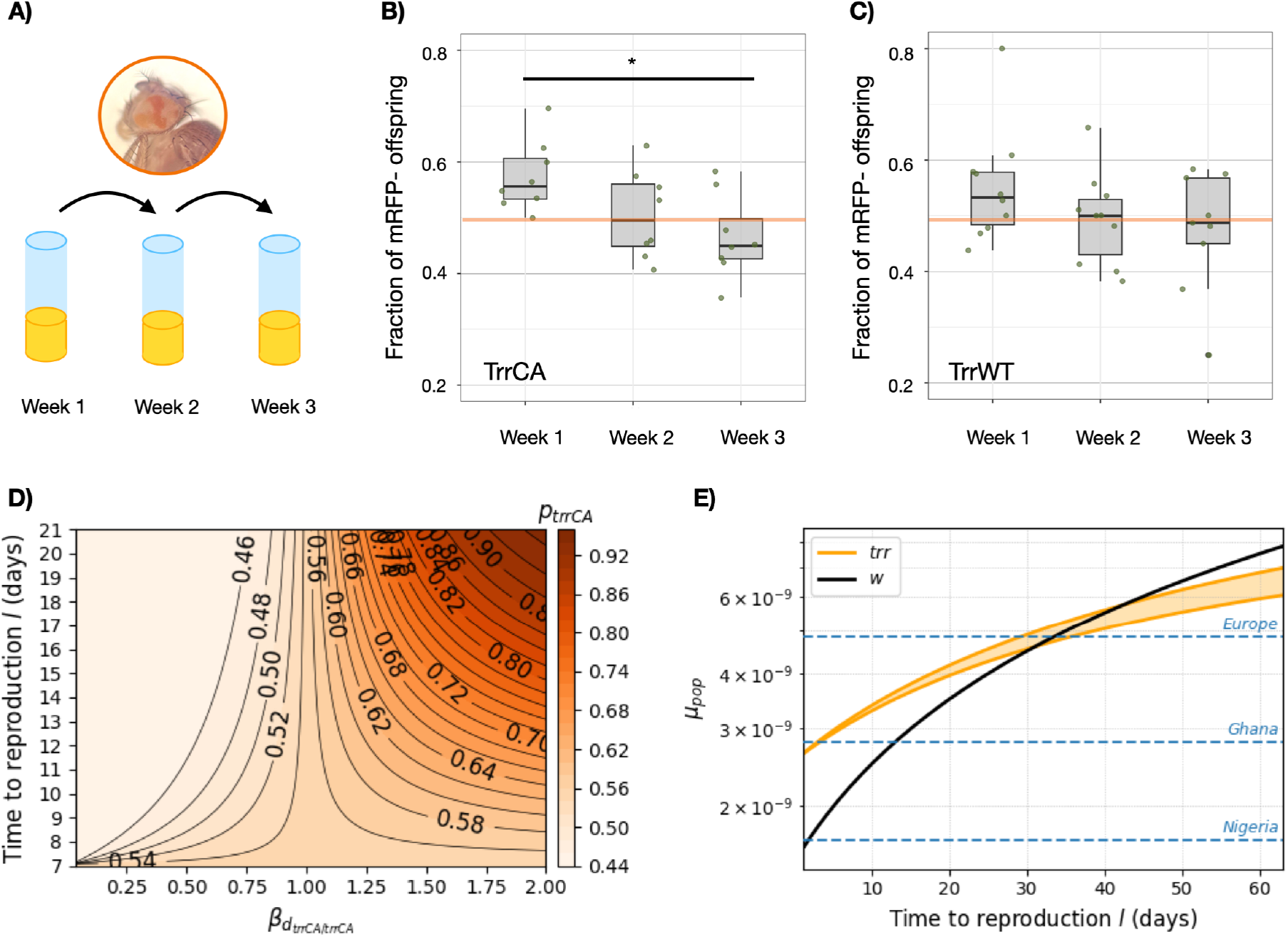
The implications of cellular fitness effect of mutations in adult female germline vary throughout individual lifetime. **(A)** Mosaic flies from the experiment illustrated in 2A were passed every week to new vials. **(B, C)** Fraction of offspring from mosaic females that did not inherit the *mRFP* marker allele (*p*_trrCA_) in each week when the alternative allele was TrrCA (B) or TrrWT carrying the *w*^1118^ variant of the *white* gene (C). Significant differences are indicated by asterisks (*p* < 0.05). **(D)** Probability of sampling the TrrCA allele upon reproduction (*p*_trrCA_) as a function of the cellular fitness effect of the mutation in homozygous cells, 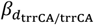, within the adult female germline of *Drosophila* and the time to reproduction, *l*, after pupation. 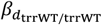 was assumed neutral relative to the backgrund heterozygous cells. **(E)** Expected generational mutation rate per locus, *µ*_*pop*_, as a function of the time to reproduction following pupation, *l*, predicted by the *Drosophila* germline model. The orange region indicates the range of mutation rates predicted for the *trr* gene given the range of cellular fitness effects estimated to reproduce the observed temporal pattern of *p*_trrCA_ in our experiment (Fig. S3). The black line indicates the predicted mutation rate for a neutral locus, corresponding to the *white* gene in our experiment (C). For these estimations the developmental parameters are the same as in Fig. 2E, but with a cellular fitness effect during development, *β*, of 1.2 in the case of *trr* and 1 in the case of *w*. The turnover rate, *d*, for the maturation phase of our models was 0.038 (Methods). Blue dashed lines indicate genome-wide mutation rates per locus estimated for *Drosophila* populations from three locations (128, 129).

Together, these results indicate that the selective regimes acting during development and adulthood can act in opposite directions, further complicating the interpretation of *µ*_*pop*_ estimates. Thus, as in the case of differences between zygotic and germline cellular fitness effects (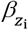 and 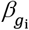 Fig. S1C) intraorganismal selection during the lifespan of a mature organism may interact with selection acting earlier in development to shape the effective 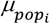 (Fig. S3B). Evaluating how the reversal of the cellular fitness effect would affect mutation rates at the *trr* locus, given the inferred range for 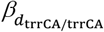 and the architecture of the *Drosophila* female germline, mutation rates at *trr* are expected to increase with age, but substantially less than those at loci where mutations are neutral within the germline, such as in the *white* gene (Fig. 3E). Thus, the reduced model-predicted mutation rates during adulthood partially offset the initially elevated mutation rates resulting from the higher cellular fitness of TrrCA during development (Fig. 3E).

Our results provide direct experimental evidence that developmental and life-history parameters can determine how cellular fitness effects translate into transmission bias and, therefore, estimates of heritable mutational input can vary even under constant molecular mutagenesis. Furthermore, the inferred reversal in the cellular fitness effect of TrrCA indicates that cellular fitness landscapes can be dynamic within individuals: the effects of cell-lineage selection on inheritance depend on the timing, magnitued and sign of selection, all of which may vary during the life cycle.

### The cellular distribution of fitness effects, genome-wide mutation rates and identity composition of heritable mutations

Our experiments with the mutants of *w* and *trr* illustrate how mutations are expected to differ in their effects on cellular fitness across genes, depending on their functional consequences. Importantly, the resulting differences in mutation rates between these genes vary substantially with life history and, depending on the time to reproduction, span the range of genome-wide mutation rates inferred for *Drosophila* populations from different locations (Fig. 3E). Because genome-wide mutation rates are aggregates across loci (e.g. (67)), such locus-specific differences can influence the genome-wide estimate when they are distributed systematically across the genome (68). Thus, genome-wide mutation rates may depend not only on the intrinsic molecular propensity of loci to mutate, but also on the cellular distribution of fitness effects across the genome (cDFE).

To explore the potential magnitude of this effect, we emulated the conventional estimation of genome-wide mutation rates by calculating *µ*_*pop*_ as the average mutation rate across loci, assigning each locus a cellular fitness effect (*β*_*i*_) drawn from a specified cDFE. To our knowledge, cDFEs have not yet been empirically characterized for any multicellular organism. We therefore considered a biologically plausible, yet illustrative, distribution in which lethal mutations account for roughly 30% (69), while non-lethal mutations are centred around neutrality with a predominance of deleterious mutations (Fig. 4A, Methods), broadly resembling distributions of fitness effects inferred from yeast cell growth assays (70, 71). We additionally carried out a sensistivity analysis exploring distributions containing a variable tail of mutations that increase cellular fitness, occurring with probability 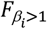 (Fig. 4A). Across a range of developmental parameters, genome-wide *µ*_*pop*_ depended strongly on the interaction between organismal traits and cell-lineage selection for all cDFEs explored (Fig. 4B). Although mutations increasing cellular fitness are generally expected to be rare (72), even a small fraction of such mutations substantially increased estimates of genome-wide *µ*_*pop*_ (Fig. 4B). Across the cDFEs considered here, these results indicate that developmental and life-history variation can interact with cell-lineage selection to produce substantial differences in generational mutation rates among species, even when the underlying cDFE is predominantly neutral or deleterious, and that the effect may be strongly affected by the frequency of mutations that increase cell fitness.

**Figure 4.**
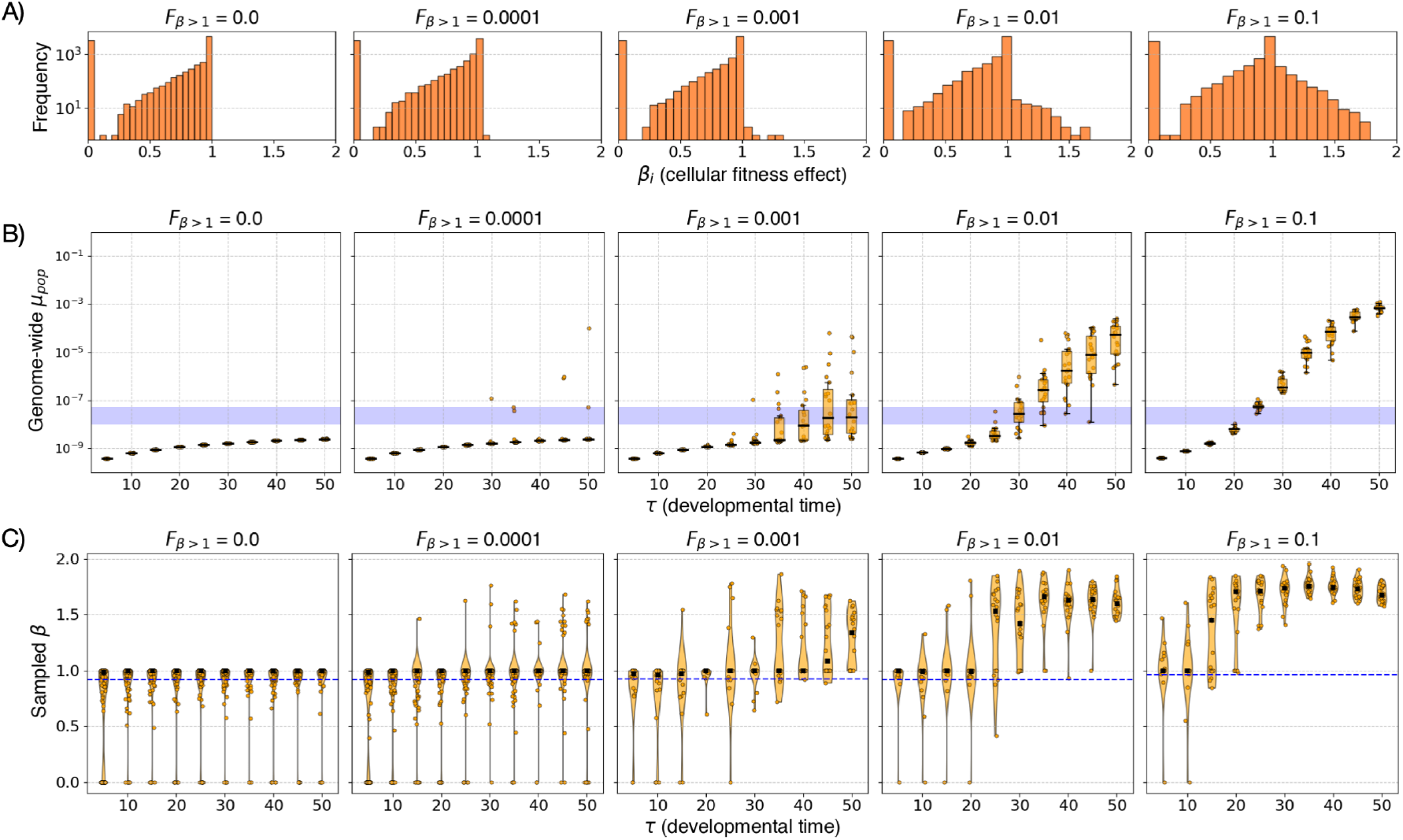
The cellular distribution of fitness effects shapes generational mutation rates and produces qualitative biases in the transmission of new genetic variants. **(A)**Distribution of cellular fitness effects (cDFE) of mutations. The shape parameters for each of these distributions correspond to the baseline described in the Methods. For each cDFE, the proportion of mutations that could increase cellular fitness during development (*β*_*i*_ > 1) is given by 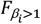, which is indicated above each panel. The proportions of deleterious and neutral were such that 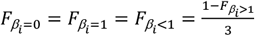. **(B)** Genome-wide *µ*_pop_ as a function of developmental time, *τ*, for different cDFEs considering the 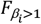 values indicated in the corresponding above panels. For each value of *τ* we calculated genome-wide *µ*_pop_ considering 20 different random cDFEs. Purple band indicates the range of empirical estimations of *µ*_pop_ for vertebrates (44) as a reference. All developmental parameters are here set to their baseline values (*β*_*I*_ = 1, *µ*_*mol*_ = 1*x*10^−10^, *C* = 1, *γ*=1, *τ* = 20), and no maturation phase was considered. **(C)**We simulated the sampling of a mutant gamete and estimated the likely value of *β*_*i*_ of the mutant locus. The blue dashed line indicates the expected value given each cDFE if there was no cell-lineage selection. Note the transmission bias towards higher sampled *β*_*i*_ as a function of the distribution of fitness effect and developmental time.

We also expected that the interaction between developmental parameters and the cDFE would influence not only the quantity of heritable variation entering populations but also its composition. Specifically, we predicted that this interaction would generate transmission biases favouring mutations with particular cellular fitness effects. To examine this effect, we simulated development and sampled 100 mutant gametes, recording the cellular fitness effect (*β*_*i*_) of each transmitted mutation. Under developmental regimes promoting stronger cell-lineage selection, transmitted mutations were significantly enriched for variants that increased cellular fitness, whereas mutations with deleterious cellular effects were underrepresented (Fig. 4C).

Together, these simulations indicate that developmental and life-history parameters, cell-lineage selection, and cDFE jointly shape not only the number of new alleles entering a population, but also the cellular fitness composition of those alleles. Thus, the cellular fitness effects of mutations may influence not only how much heritable variation enters populations, but also which classes of mutations are preferentially transmitted.

## Discussion

### Germline cell-lineage selection and the causes of mutation rate variation

Mutation rates are generally viewed as properties of molecular mutational processes, shaped by factors such as DNA replication frequency and fidelity, as well as the balance bewteen DNA damage and repair. Here, we emphasize a view of them as emergent properties of organismal biology (32) while also accounting for the cellular fitness effects of mutations within organisms. By integrating mathematical models with experimental evidence, we demonstrate that selection among cell lineages interacts with developmental architecture and life-history traits to prodcue specific transmission biases, which repercusses on both the rate at which new mutations enter populations and the composition of those mutations. This perspective provides an extended framework for understanding how developmental and life-history evolution may contribute to mutation-rate variation. Because the strength and direction of cell-lineage selection depend on organismal traits, evolutionary changes in development, germline organisation, and longevity have the potential to alter not only the quantity of heritable variation produced each generation but also the classes of mutations that are preferentially transmitted.

Because each locus may have its own selective interplay with the organismal environemnt, our results reinforce the view that locus-specific features must be considered when interpreting genome-wide mutation rates (68), which are also known to depend on factors such as nucleotide composition (73), transcriptional activity (74) and epigenetic marks (75, 76). Furthermore, because the influence of cell-lineage selection depends on traits that vary from species to species such as developmental duration, rates of cell division, time to reproduction or, potentially, the cellular distribution of fitness effects, comparisons of mutation rates and rates of molecular evolution between species such as mice and humans (46), or across taxa as disparate as shrews and orcas, or snowy owls and pelicans (44, 77), should be interpreted with considerations of the effects of cellular-level selection in addition to differences in molecular mechanisms and life-history strategies.

The generation time effect provides a useful example. This long-recognised pattern describes the tendency for species with longer generation times to exhibit lower rates of molecular evolution (46, 47), which may be attributed to a negative relationship between the yearly mutation rates of species and their generation times (44, 78). While reporting this trend, Lewin and Eyre-Walker (78) recently discussed several explanations, including elevated mutagenesis during early development and prenatal life (79, 80), which could cause short-lived species to accumulate more mutations per unit time; higher germline cell-division rates in short-lived species, as previously proposed (81); and differences in DNA damage, repair, and purifying selection against mutator alleles consistent with the drift-barrier hypothesis. Despite their differences, these explanations all treat mutation-rate variation principally as a consequence of changes in the production of mutations. Within the framework we are here proposing, we can consider a complementary – or alternative-cause based on the selective filtering of mutations after they arise. In our model, mutant and wild-type germline stem-cell lineages continue to compete throughout the reproductive lifespan of an individual. Because longer-lived organisms experience this selective process over a longer period before reproduction, mutations with deleterious effects on cellular fitness become progressively underrepresented in the gamete pool, whereas mutations that increase cellular fitness become enriched. Under biologically plausible cellular distributions of fitness effects (cDFEs) dominated by deleterious or nearly neutral mutations, as observed in yeast growth assays (70, 71), this mechanism can generate a negative relationship between yearly mutation rate and generation time (Fig. S4, S5), consistent with empirical trends (78). Germline cell-lineage selection may therefore represent an additional evolutionary process contributing to the generation-time effect by reducing yearly mutation rates without requiring any change in the underlying molecular mutation process. However, the validity of this hypothesis ultimately relies on the true cDFEs across species, which, as we argue below, remain an open empirical question.

### Evolutionary implications of germline cell-lineage selection

If cell-lineage selection is an important determinant of genome-wide mutation rates and transmission biases, it may also represent a significant evolutionary force. By acting on new mutations before transmission, cell-lineage selection can provide a relatively “cheap” way to purge deleterious variants (17, 18, 20), it may affect the probability of fixation of variants that alter cell fitness (18), facilitate the recurrent appearance of alleles that are deleterious at the organismal level (20, 27-29, 31)— as has recently been suggested for the persistence of disease-causing variants in human populations (30) —and promote evolvability when the effects of mutations on cellular and organismal fitness are aligned (19). Crucially, the cellular fitness effects of mutations are locus-dependent, and, as we have shown, cell-lineage selection may therefore bias both the composition of allelic diversity and the diversification rates of each locus within populations. We further propose that, because biases in mutational availability can shape the set of adaptive paths open to evolution (82-87), cell-lineage selection may consequently influence which evolutionary trajectories are more likely to be followed by populations and species. An immediate implication of this is that cell-lineage selection may influence the regions of genotype space that populations explore over evolutionary time (Fig. 5), and that selection at the cellular level affects not only the amount of heritable variation entering populations and the rates of diversification, but also the direction of evolutionary change.

**Figure 5.**
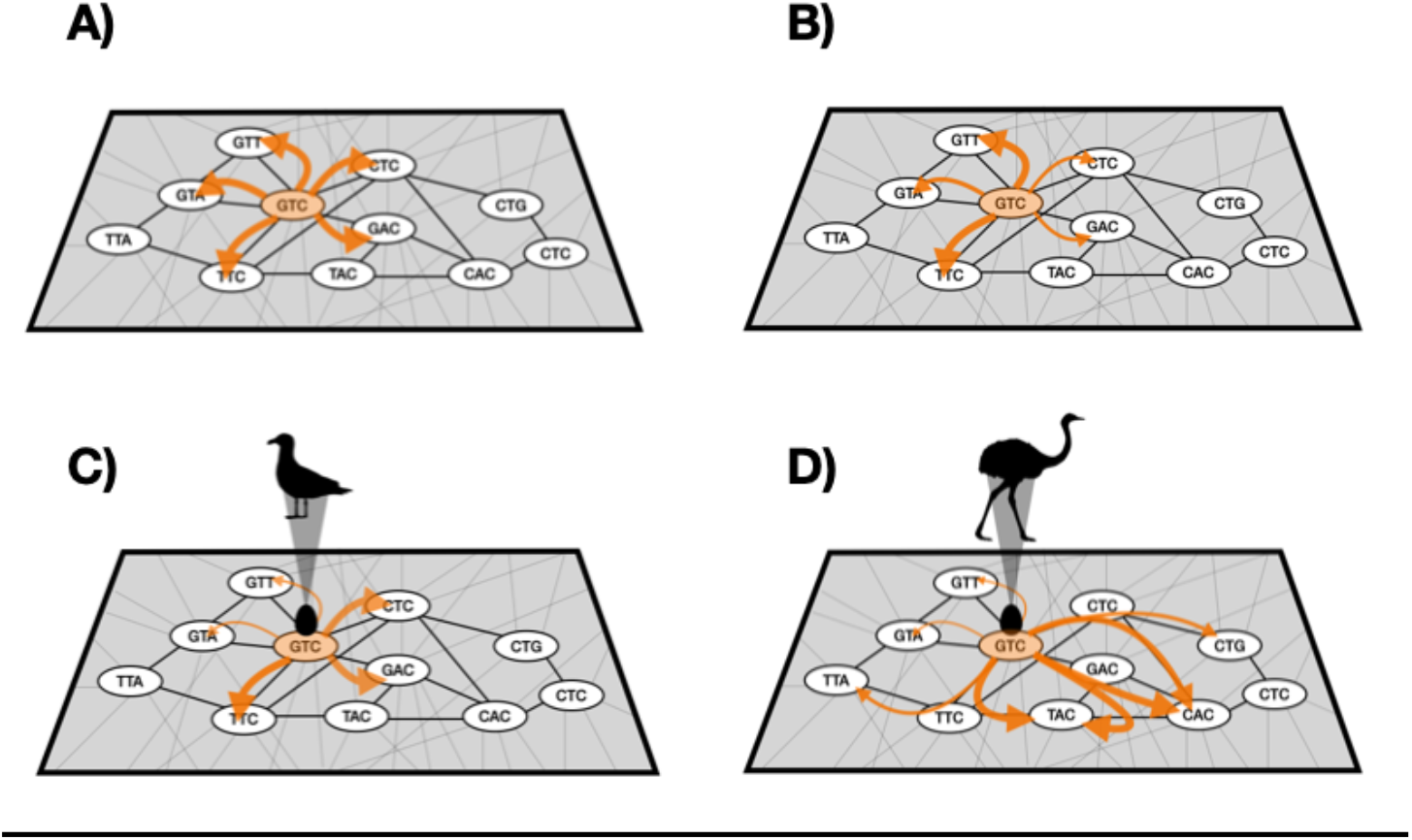
How development can influence the trajectories and pace of genotypic evolution. In a theoretical model of evolution, it could be considered that given a starting genotype, GTC, all mutational steps are equally likely (A). However, if molecular realism is taken into account there are certain mutations that are more likely than others, as represented by arrows of different thickness going to CTC and GAC. (B). Furthermore, as we argue in this work, development can act as a selective filter for mutations which increases or decreases the likelihood of mutational steps depending on how a mutation affects the cell lineage it belongs to (C). Developmental and life-history differences can also impact how far mutational steps can take an organism (D).

Importantly, the different evolutionary consequences of cell-lineage selection are themselves expected to depend on developmental and life-history evolution. Because traits such as body size, developmental duration, rates of cell division, and generation time evolve among lineages, the importance of cell-lineage selection in shaping evolutionary trajectories may also evolve. This means that, in the light of cell-lineage selection, the molecular evolvability of Darwin’s rhea is not the same as for a seagull (Fig. 5) – large-bodied, long-lived species are expected to differ from small-bodied, short-lived species not only in the frequency with which new mutations enter populations (32, 88), but also in the duration and intensity of the intraorganismal selective processes. In this view, the evolutionary consequences of cell-lineage selection become contingent on the organismal traits that define the selective environment experienced by germline cell lineages. Thus, under this framework genotype-phenotype maps play a role in evolution that goes beyond structuring the genetic availability of phenotypic variation (89-91); they also allow phenotypes to directly influence how genetic variation is produced.

Variation in developmental and life-history traits occurs not only among species but also within populations. Because individuals differ in these traits, the effects of cell-lineage selection on mutation rates and the directionality of mutational change are likely to vary among individuals. Such variation may involve quantitative traits, including body size and age at reproduction. Notably, it may also be represented by sexually dimorphic traits, particularly germline architectures. In mammals, for example, females show little or no turnover of gamete-capable cells after maturation (*d* = 0 in our model), whereas males maintain substantial post-maturational turnover (*d* > 0), implying more prevalent cell-lineage selection on paternally derived mutations (Fig. 1D, S1D). Within a population, this could substantially bias not only the number of paternally inherited mutations but also their composition, enriching the paternal mutational input for mutations that increase cellular fitness.

Additionally, we expect cDFE to be, like development and life history, an evolvable feature that helps determine how cell-lineage selection shapes the rate and direction of genetic diversification within species. Once a mutation affecting cellular fitness arises, the intraorganismal selective landscape experienced by cells carrying subsequent mutations changes. Mutations that were previously advantageous at the cellular level may therefore no longer be favored in the new genetic background. This process could represent a form of diminishing-returns epistasis, in which the fitness effect of a mutation is smaller in a fit genotype than in an unfit genotype (92-94). If cDFE so evolves, then it is likely that mutation rates and transmisison biases would evolve alongside it, as each cDFE interacts with organismal features to shape the rates and identity of heritable mutations (Fig. 4). This perspective is supported by studies showing how in mutation accumulation experiments in *Drosophila* lineages that were loaded with deleterious variants tended to have higher mutation rates than wild-type lineages (95, 96), and also why during mutation accumulation there tends to be a decrease in mutation rates over generations (97). It is also consistent with recent evidence that the proliferative advantage conferred by oncogenic mutations can depend strongly on genetic background in mice (98).

### The prevalence and relevance of germline cell-lineage selection

The importance of cell-lineage selection in shaping population genetic diversity ultimately depends on how prevalent it is within the germline of organisms, which is still an empirically underexplored question. Our experiments in *Drosophila*, together with previous experimental (22, 23, 27), theoretical (17-19, 21), and sequencing (30) studies, do support the view that cell-lineage selection can meaningfully influence inheritance and mutation-rate variation. However, these studies are mainly concerned with limited sets of mutations, genomic regions, and experimental systems. Evaluating the impact of cell-lineage selection on genetic evolution requires a deeper understanding of its selective dynamics in its organismal context and, crucially, the cDFE. Genome-wide cDFE are still far from being known. Even the most comprehensive studies to date evaluating selection in the human germline (30) focus largely on subsets of genes and coding regions. To properly characterise cDFE, further studies should go beyond the examination of coding regions, since variants in non-coding regulatory elements such as promoters and enhancers can alter gene activity and cell proliferation, as shown for regulatory variants affecting the proto-oncogene *myc* (99). Emerging approaches that combine deep sequencing with barcode-guided cell-lineage tracing (100) and phylodynamic reconstruction of developmental cell lineages (101) are likely to provide the empirical data needed to estimate cDFEs and evaluate the evolutionary importance of cell-lineage selection across different organsimal contexts.

Furthermore, exploring the evolutionary consequences of cell-lineage selection for species evolvability will require integrating models of organismal evolution that incorporate cell-lineage selection with population-genetic models of adaptation. Our results represent a first step in this direction by incorporating cell-lineage selection into developmental models of mutation-rate evolution. The framework nevertheless relies on several simplifying assumptions, including the omission of ploidy, a simplified representation of developmental architecture, and the assumption that cellular and organismal fitness are largely decoupled. Although these assumptions may be of relevance, they do not undermine the model, recent results have shown that the efficacy of ploidy in masking deleterious mutations may be weaker than anticipated (102), evidence for germline cell-lineage selection has now been reported across organisms with markedly different germline architectures, including plants, insects, and mammals (21-23, 30, 103), and abundant clonal expansions in healthy tissues indicate that cell-level selection is a pervasive feature of multicellular biology rather than necessarily a pathological process reducing organismal fitness (10, 104-107). More biologically realistic models will be required to weight the relevance of cell-lineage selection for the genetic diversity in populations, but ultimately the prevalence and evolutionary importance of cell-lineage selection remain empirical questions. Addressing these questions will require a much deeper characterization of genome-wide cDFEs across multiple layers of cellular physiology and of their interactions with neighboring cells and whole-organism physiology.

### Outlook

Overall, our results suggest that mutation rates are best understood not as fixed molecular constants, but as emergent properties arising from interactions among molecular mutagenesis, developmental architecture, life-history traits, and selection among cell lineages. Together, these factors can influence both the tempo and the direction of evolutionary change, implying that the evolution of developmental and life-history traits may also modify the mutational and selective processes that shape population-genetic dynamics. Cell-lineage selection thus provides a window into the broader question of how organismal organization contributes to shaping evolutionary trajectories (108), emphasizing how evolutionary trends, instead of being solely the product of chance and an externally imposed necessity, are also actively shaped by the internal properties and predispositions of organisms themselves.

## Methods

### Developmental model for estimation of *µ*_pop_

We define the generational mutation rate at locus *i*, 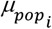, as the probability of sampling a mutant cell among all cells in an organism that are capable of producing gametes at the time of reproduction of an individual. To calculate this probability, it is necessary to first calculate *M*_*i*_, which represents the total number of cells that are expected to carry a mutation at a given locus *i*. In previous work (32), we adapted a model wherein *M*_*i*_ is a function of the molecular mutation rate *µ*_*mol*_, the total developmental time *τ*, the rate of cell divisions during development *C* and the time to germline sequestration *γ*. Following Otto & Orive (18), we now added a factor *β*_i_, which represents the effect of a mutation on the fitness of a cell carrying a mutation on locus *i*. Cellular fitness could be affected by mutations involved in various processes, including the migration of cells, proliferation, and, in the specific case of germline cells, the determination and maintenance of the germline cell fate. Because of these effects, such mutations may end up under- or over-represented in the gamete pool produced by an organism (28, 109). In our model, as in (18), *β*_*i*_ specifically refers to how changes in the rates of proliferation during development may influence the representation of mutations at locus *i*. Deleterious mutations are not necessarily lethal to cells, nor do they imply a complete cessation of their proliferation. Instead they could simply make cells proliferate slower (e.g. (110)). On the other hand, beneficial mutations would be those that increase the rates of cell division (e.g. (111)).

When considering mutations to be of replicative origin (i.e. the units of *µ*_*mol*_ are per cell division), at cellular generation *x*, one would expect 2 ^*x*^ (1 − *µ*_*mol*_) ^*x−1*^ cells to not carry any mutations. Each of those cells can mutate with a probability given by *µ*_mol_ and the mutated lineage will then undergo *β* _*i*_(*C τ - x*) cell divisions more until the conclusion of development. Thus, the number of mutant cells resulting from mutations at time *x* is:

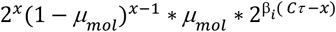

Summing over all *x* gives us *M*_*i*_ at the conclusion of development:

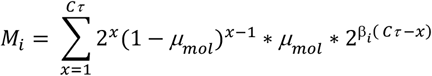

For estimating *µ*_pop_ it is also necessary to calculate the expected number of cells that do not undergo mutations at locus *I* through the course of development i.e., *N*_i_. This can be calculated as:

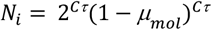

The total number of cells at the end of development is (*M*_*i*_ + *N*_*i*_) and thus 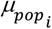 approximates the probability of sampling a cell mutated at locus *i* from all cells in the pool of gametes, such that

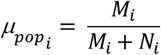

An additional relevant developmental parameter is the timing of germline sequestration, *γ*, which varies widely across the tree of life (112, 113), even between closely related species (114). This parameter can structure the germline cell lineage in a way that can influence the selective dynamics at the cellular level (115, 116), and therefore, its impact on the proportions of mutant cells at the end of development. When considering our germline sequestration parameter, *γ*, we fractioned our calculation of *M*_i_ and *N*_i_ as zygotic (*M*_z_, *N*_z_) and germline (*M*_g_, *N*_g_) components, such that:

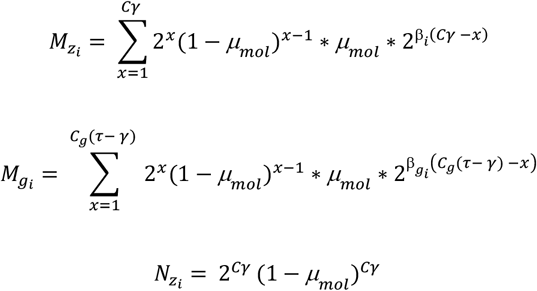

and

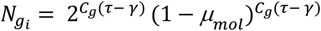

Where *C*_g_ is the rate of cell divisions in the germline cell lineage after its sequestration and 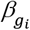 is the cellular fitness effect on a mutation specifcially in the germline lineage. We thus calculated the probability of sampling a cell that mutated by the time of germline sequestration, 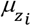, as,

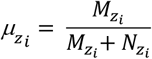

and the probability of sampling a mutated gamete from a mutation that arose after germline sequestration, 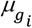, as,

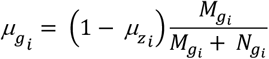

And,

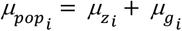

For a value of *τ* = 20, we explored a range going from *γ* = 1, to *γ* = 20. This range encompasses scenarios from very early germline sequestration, exemplified by *Drosophila*, in which sequestration occurs after roughly 9–10 cell divisions (117), to very late germline delineation, traditionally—though not without dispute (118)—thought to occur in plants.

The calculations shown above correspond to those considering *µ*_*mol*_ to track cell divisions. In the cases in which mutation rates are taken to track time rather than cell divisions, which result mainly from the contribution of the balance of DNA damage and repair (43, 55, 119, 120), the calculations for *M*_*i*_ and *N*_*i*_ were modified such that the sum is taken over each time unit *t* that go from the start of development *t = 1*, to its end *t* = *τ*. In this case the number of mutant cells at the end of development that arise at time *t* are given by the product of the cells that have not mutated thus far but do mutate at this point, (2^*ct*^(1 − *µ*_*mol*_)^t−1^ * *µ*_*mol*_ ), and their clonal expansion for the rest of development 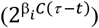, such that,

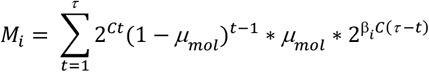

And,

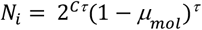

Note that in this case the probability of not having mutated throughout development (1 − *µ*_*mol*_)^*τ*^ is not a function of the rate of cell divisions, but only *τ*.

### Expansion of the model to account for maturation and time to reproduction

To consider the role of lifetime of an organism in the calculation of 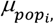 we considered the pool of cells resulting from the developmental process to be constant in number. Immediately at the end of development, there are *M*_*t=*0_ cells in the germline stem cell niche that are mutated and *N*_*t=*0_ that are not. During an individual’s lifetime, germline stem cell niches can undergo a cellular turnover. In mice, for example, spermatogenic stem cells sometimes cease to be stem cells and are replaced by neighboring stem cells, keeping the number of stem cells relatively constant (121). Although the repopulation of these cleared niches by neighboring cells could be a stochastic process (122), mutations that affect the rate of stemness loss (either through death or differentiation) might eventually create a mosaic of spermatic stem cells with an overall higher fitness (28, 109). Thus, a selective cell turnover might impact the repertoire of mutations that a single individual can introduce into a population’s gene pool.

We considered a time to reproduction *l*, which is the length between the end of development and reproduction, during which germline stem cells can undergo a turnover (due to differentiation or death) at a rate *d* over an arbitrary unit of time *t*. The rate at which mutant cells replace lost cells during this turnover is given by a selective variable *β*_*l*_. As in the case of the developmental selective parameter *β*, when *β*_*l*_ < 1 mutations are deleterious, when *β*_*l*_ = 1 mutations are neutral and when *β*_*l*_ > 1 mutations are advantageous to cells. In our model, during the lifetime of an organism before reproduction, it is possible for the total pool of mutated cells to decrease either due to reverse mutations (which would occur at a low rate, *M*_*t−1*_*µ*_*mol*_) or loss due to death or differentiation (*dM*_*t−1*_). Furthermore, mutant cells can increase in proportion whenever non-mutant cells in the stem cell niche mutate (*N*_*t−1*_*µ*_*mol*_ ) or whenever mutant cells proliferate 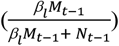 to replenish the stem cell niche after the loss of sibling cells (*d*(*M*_*t−1*_+ *N*_*t−1*_)). The number of mutated cells at time *t* during the lifetime of an organism can thus be calculated as,

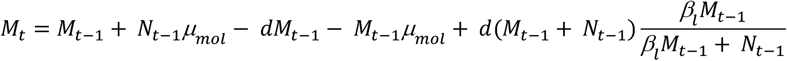

And the number of non-mutant cells would be given by

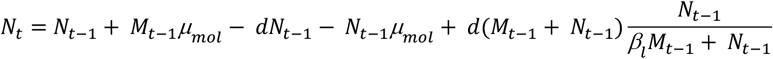

Following this logic, we can also consider the effects of mutations that affect the rate of loss of cells in the germline niche, *β*_*d*_, such that

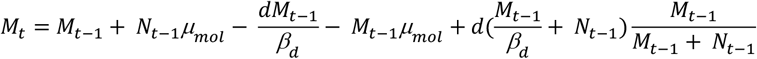

And the number of non-mutant cells would be given by

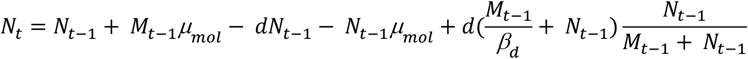

### Genome-wide generational mutation rates

To estimate *µ*_pop_ across the entire length of a genome, as a first order approximation, we assumed that no there is no interaction between the cellular fitness effects of different mutations. Using this simplification, we can estimate *µ*_pop_ across the entire length of a genome by averaging *µ*_*pop*_ across all loci which may have different values of *β*_i_.

For this, we modeled the cellular distribution of fitness effects (cDFE) using a five-component mixture distribution representing discrete and continuous classes of *β*_i_. In this framework, fitness effects are encoded on a continuous scale where values of 0 correspond to strictly deleterious mutations, 1 to strictly neutral mutations, and 2 to highly beneficial mutations. The mixture model comprises point masses at these three canonical fitness classes (0, 1, and 2), together with two continuous components capturing variation within deleterious and beneficial effects. Specifically, deleterious effects were modeled using a Beta distribution with shape parameters *a*_;_ and *b*_;_, yielding values between 0 and 1. Beneficial effects were modeled using a Beta distribution with shape parameters *a*_<_ and *b*_<_, shifted by +1 to yield values between 1 and 2. The relative contribution of each component was governed by mixture weights 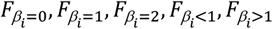, which were normalized to sum to one.

To estmate genome-wide *µ*_pop_ we considered our model for a total of 10000 loci for which *β*_i_ corresponded to different cDFE generated such that the shape parameters governing the continuous components of the cDFE were fixed at *a*_;_ = 2, *b*_;_ = 0.6 for the deleterious Beta distribution, and *a*_<_ = 0.6, *b*_<_ = 4 for the beneficial Beta distribution. The weight of the intermediate point-mass component was set to 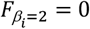. To investigate the effect of beneficial mutations, we varied the mixture weight 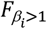 across the set {0, 10^-4^, 10^-3^, 10^-2^, 10^-1^}. For each value of 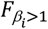, the remaining mixture weights were adjusted symmetrically such that 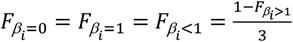, ensuring that all mixture weights summed to one. For simplicity, we only accounted for a developmental phase in these calculations with fixed parameters as in their baseline with *µ*_*mol = 1*_*x*10^−10^, *C* = 1 and *γ* = 1. For each set of developmental and cDFE paramters we considered 20 different cDFEs for 10000 loci to estimate genome-wide *µ*_pop_ as the mean *µ*_pop_ of all loci. Using these same cDFEs, we calculated the *µ*_*pop*_ associated with each sampled *β* value. Mutations were then probabilistically transmitted according to these values using Bernoulli sampling, allowing us to estimate which classes of cellular fitness effects are enriched among transmitted mutations. For each parameter combination, simulations were run until 100 transmitted mutations were obtained.

### Flippase assay to assess cell competition in fly germline

We developed heat shock-induced mitotic clones in fruit flies following a standard protocol (66). Briefly, a fly line (# 31418 from the Bloomington Drosophila Stock Center) containing a red fluorescent protein (mRFP), a flippase-encoding gene induced by a heat-shock promoter (hsFLP) and a FRT site for the binding of flippases was crossed with another line carrying a single-amino acid substitution in the *trithorax-related* gene (*trr*) and the *w*^1118^ allele for the gene white. We thus created a *trr*^CA^,*w*^1118^,FRT,hsFLP and a *w*^1118^,FRT,hsFLP lines. For our experiments we crossed these lines with the parental mRFP,FRT,hsFLP lines and we exposed the heterozygotic embryos collected during a window of four hours to a heat shock of 37°C for one hour.

We collected mosaic virgin females that developed from these heat-shocked embryos as well as virgin females from embryos that were not heat-shocked (control). Each of these virgins were individually paired to two *w*^1118^ males and we then counted the fraction of offspring that were positive for red fluorescence, which are the ones that inherited the mRFP+ allele and not the mRFP- (*trr*^CA^,*w*^1118^) allele. For the same set of pairs, we evaluated the fraction of mRFP+ pupae from offspring produced during the first week immediately after mothers were collected, as well as during the second and third weeks.

### Estimation of *β* and *β*_*l*_ in the female germline of *Drosophila*

To adapt our model to *Drosophila* female germline development and match it with our experiments, we considered developmental stages during which the germline population undergoes appreciable proliferation. Briefly, during early embryogenesis, the embryo develops as a syncytium, and a approximately 10-15 nuclei migrate to the posterior germ plasm, where they become cellularized to form the primordial germline, or pole cells (123). These cells undergo up to two additional rounds of mitotic divisions, generating a population of approximately 30-40 primordial germ cells (PGCs) (124). Following migration and coalescence with the somatic gonadal precursors, approximately 7-13 PGCs are incorporated into each developing gonad. During larval development, these PGCs undergo continued proliferation, reaching a population of more than 100 germ cells per gonad by middle third instar (125).

In our flippase assay we induced mitotic recombination during early development, spanning the phase of proliferation of PGCs. Consistent with what is known of *Drosophila* female germline development, we considered an initial population of *k* = 10 primordial germ cells. All of these cells would have the zygotic genotype *TrrCA*/*mRFP*. From this population, we assumed that one dividing cell would successfully undergo mitotic recombination upon induction of the flippase, such that the cellular composition at the end of this stage would then be 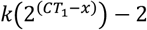 cells with the zygotic genotype (*N*_*trrCA/trrCA*_), 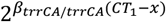 would be homozygous for the TrrCA (*M*_*TrrCA*_) and 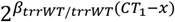 would be homozygotic for the TrrWT allele marked with mRFP (*M*_*mRFP*_). *CT*_1_ corresponds to the expected rounds of cell division during this phase (*CT*_1_ = 2, (124)), *x* represents the timepoint between 0 and *CT*_1_ at which recombination is induced, *β*_*trrCA/trrCA*_ and *β*_*trrWT/trrWT*_ represent the proliferative fitness of the respective homozygous cells relative to the heterozygotic ones. To calculate the total number of cells of each genotype at the end of development we considered how only a subset, corresponding to roughly half of the total pool of PGCs then coalesce into the forming gonad and that those cells eventually undergo *CT*_2_ rounds of cell division during larval phase (*CT*_2_ = 3.5, since the population goes from roughly 10 cells per gonad to roughly one hundred in the larva). Thus, the proportion of each genotype at the end of development is given by:

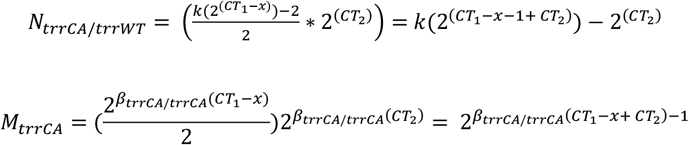

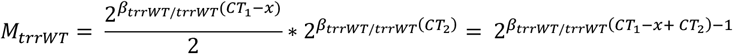

We then explored what is the expected proportion of TrrCA or mRFP+ alleles that would be sampled upon reproduction as

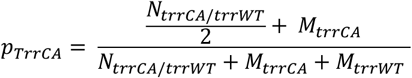

We note that this assumes that TrrCA exclusively affects rates of cell proliferation and also that to assess the effect of dominance in the fitness effects of heterozygotes it is sufficient to explore the fitness effect of TrrWT. We considered the scenario in which TrrWT is dominant in silencing the fitness effects of TrrCA in heterozygotes, but also the case in which dominance is additive such that 1 − *β*_*trrWT/trrWT*_ = *β*_*trrCA/trrCA*_ − 1.

To consider the effect of time to reproduction following development on *p*_*TrrCA*_, we considered the turnover rate, *d*, in our model. In *Drosophila*, the total number of germline stem cells decreases with age. Pan et al (126) reported a drop of around 30% from week 1 (7 days) to week 9 (63 days) in a control wild-type fly line. We thus considered the rate of stemness loss as 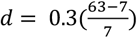. Because each ovariole only has 2-3 germline stem cells, it is unlikely that the mutational terms or turnover dynamic in our equations play a meaningful role in our experimental setup. We therefore only considered the stemness loss to estimate the cellular fitness effect of TrrCA in the female germline, 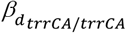, such that *p*_*trrCA*_ at time *t* would depend on

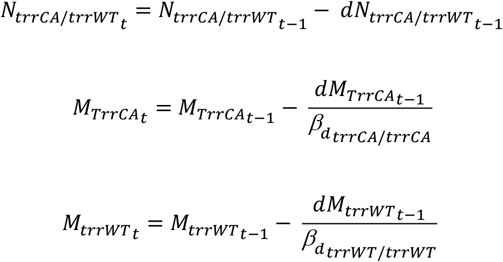

## Figures

**Table S1.**
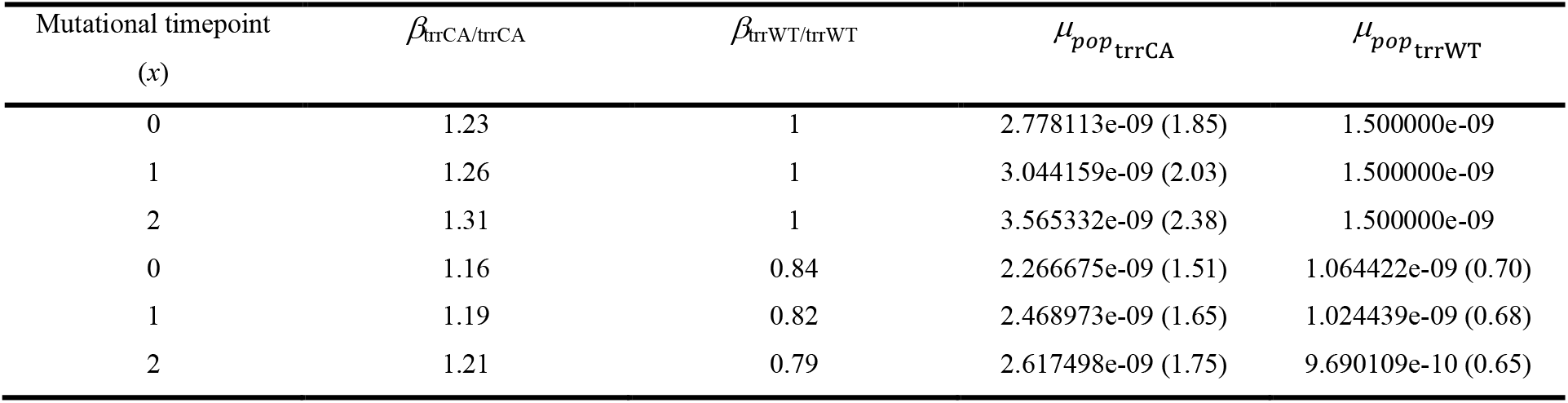
Estimates of cellular fitness effects and expected generational mutation rates of recombinant genotypes homozygous for TrrCA *and* TrrWT relative to the heterozygous state. Estimates of *β*_trrCA/trrCA_ and *β*_trrWT/trrWT_ assuming a *Drosophila* female germline architecture in which recombination occurs at time point *x* during two rounds of primordial germ cell division in early development. Using our baseline model, we also estimated the generational mutation rates at this locus corresponding to the *β*_trrCA/trrC_ and *β*_trrWT/trrWT_ values, 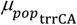 and 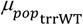,, respectively. Values in parentheses in the mutation-rate columns indicate the fold change relative to a neutral locus.

| Mutational timepoint (*x*) | *β*<sub>trrCA/trrCA</sub> | *β*<sub>trrWT/trrWT</sub> | *μ*<sub>*pop*<sub>trrCA</sub></sub> | *μ*<sub>*pop*<sub>trrWT</sub></sub> |
|---|---|---|---|---|
| 0 | 1.23 | 1 | 2.778113e-09 (1.85) | 1.500000e-09 |
| 1 | 1.26 | 1 | 3.044159e-09 (2.03) | 1.500000e-09 |
| 2 | 1.31 | 1 | 3.565332e-09 (2.38) | 1.500000e-09 |
| 0 | 1.16 | 0.84 | 2.266675e-09 (1.51) | 1.064422e-09 (0.70) |
| 1 | 1.19 | 0.82 | 2.468973e-09 (1.65) | 1.024439e-09 (0.68) |
| 2 | 1.21 | 0.79 | 2.617498e-09 (1.75) | 9.690109e-10 (0.65) |

**Figure S1.**
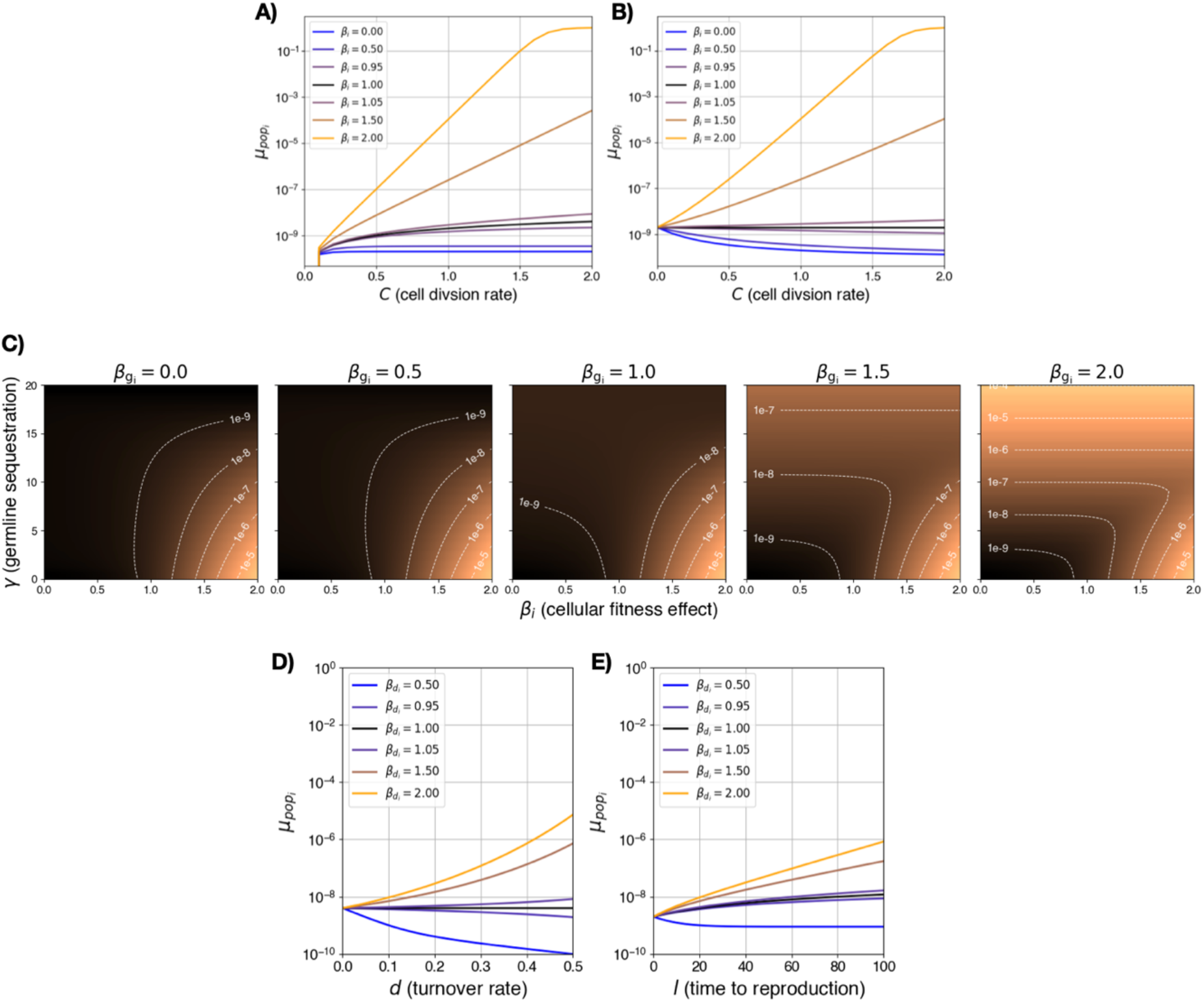
Effect on generational mutation rates of the interaction of cell-lineage selection with developmental and life history parameters under different assumptions. **(A, B) Mutations of non-replicative origin:** 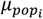 as a function of *C* for different values of the fitness effect of a mutation on locus i. *µ*_mol_ was held constant with a value of 1×10^-10^ mutations per locus per cell division (A) or 1×10^-10^ mutations per locus per unit of developmental time (B). Developmental time was held constant, such that *τ* = 20. **(C) Different cellular fitness effect before and after germline sequestration:** 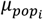 as a function of the fitness effect of a mutation on locus *i* before (*β*_*i*_) and after 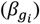 germline sequestration and the timing of germline sequestration itself. **(D, E) A cellular fitness effect of mutation affecting germline stem cell loss:** (D) *μ*_pop,i_ as a function of the rate of germline stem cell turnover (*d*) for different values of *β*_*d*_. Here, time to reproduction (*l*) was held constant at l = 10, *γ* was held at 0 and all other parameters were the same as in Fig. 1C. (E) *μ*_pop,i_ as a function of time to reproduction (*l*) for different values of 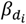. *d* was held at 0.1 and all other parameters were the same as in D.

**Figure S2.**
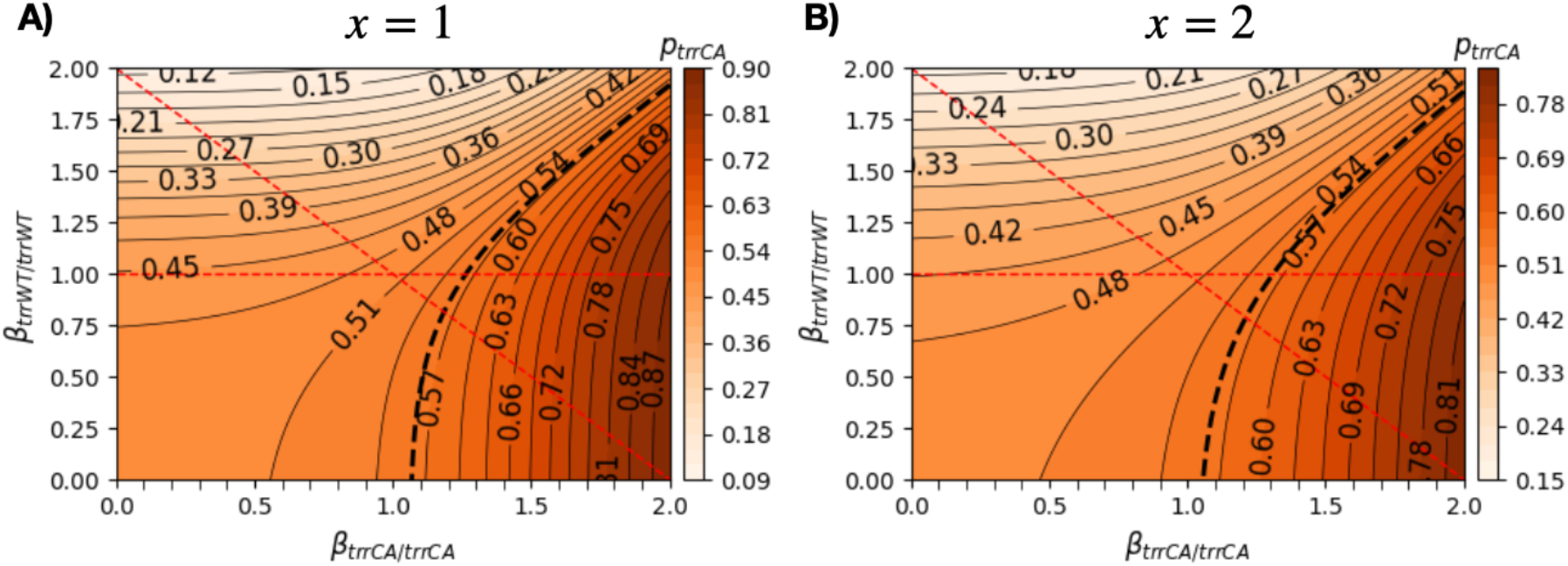
*p*_trrCA_ depending on the different possible values of the cellular fitness effect of TrrCA and TrrWT in homozigosity, *β*_trrCA/trrCA and_ *β*_trrWT/trrWT_, respecitvely. In this case *x* = 1 (A) or 2 (B), but see figure 2D for the case in which *x* = 0. The thick dashed black cline indicates the emprical expectation of *p*_trrCA_ given the results in (B). The horizontal red dashed lines represents the case in which *β*_trrWT/trrWT_ is neutral relative to the backgrund heterozygous cells, as in C, while the diagonal red dashed line indicates the case when the cellular fitness effects of the TrrCA allele are additive.

**Figure S3.**
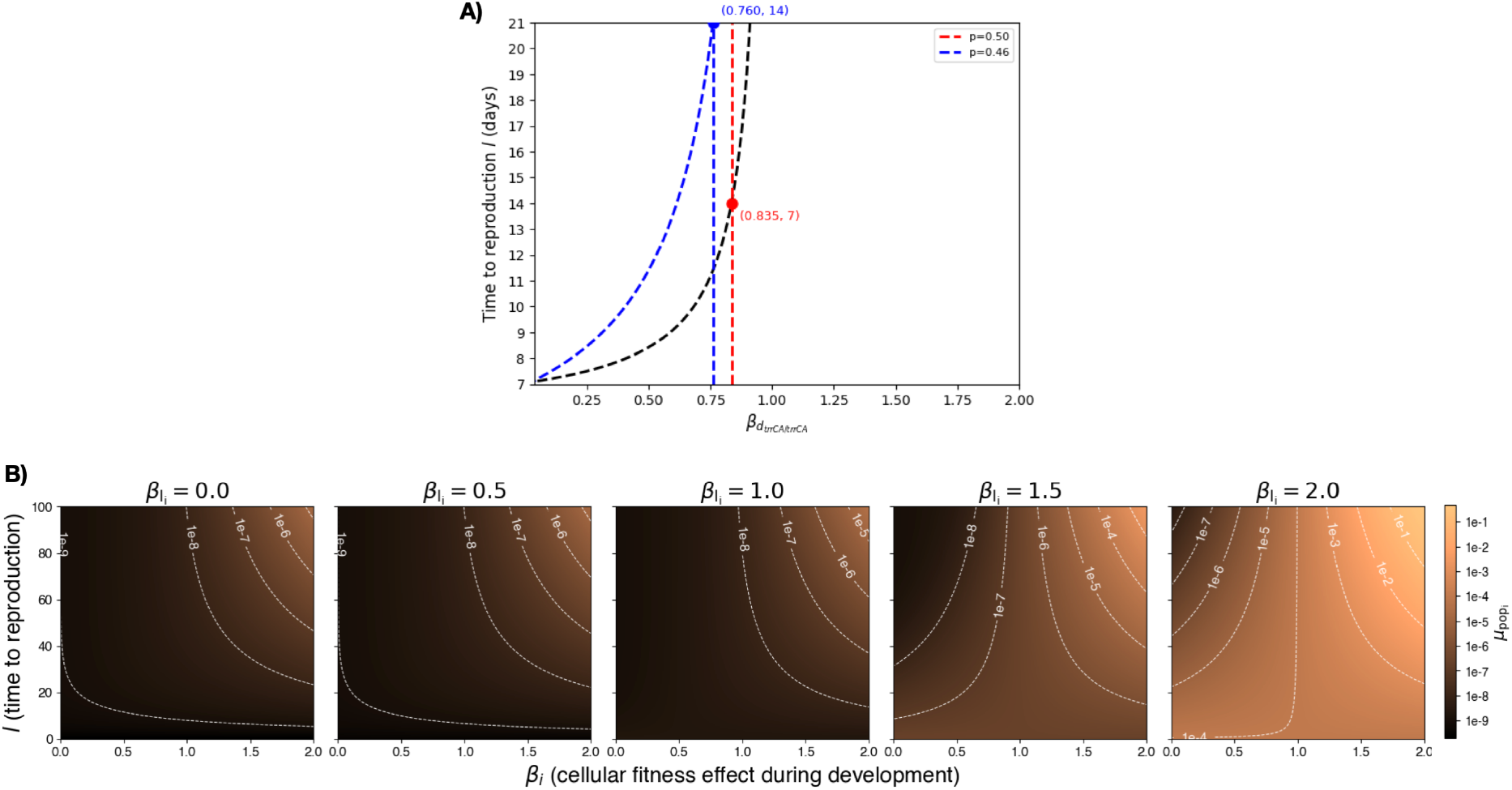
**(A)** Estimation of the range of 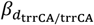 at which we expect a pattern in *p*_trrCA_ similar to the observed empirically, such that for *p*_trrCA_ at week 1 is 0.56, at week 2 (day 14) *p*_trrCA_ = 0.5 (red line), and at week 3 (day 21), *p*_trrCA_ = 0.46 (blue line). **(B)** 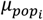 as a function of the fitness effect of a mutation on locus i before (*β*_*i*_) and after 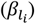 the end of development and time to reproduction *l*. All developmental parameters are here set to their baseline values (*β*_*i*_ = 1, *µ*_*mol*_ = 1*x*10^−^10, *C* = 1, *γ*=1, *τ* = 20), while stem cell turnover, *d*, was set to *d* = 0.1.

**Figure S4.**
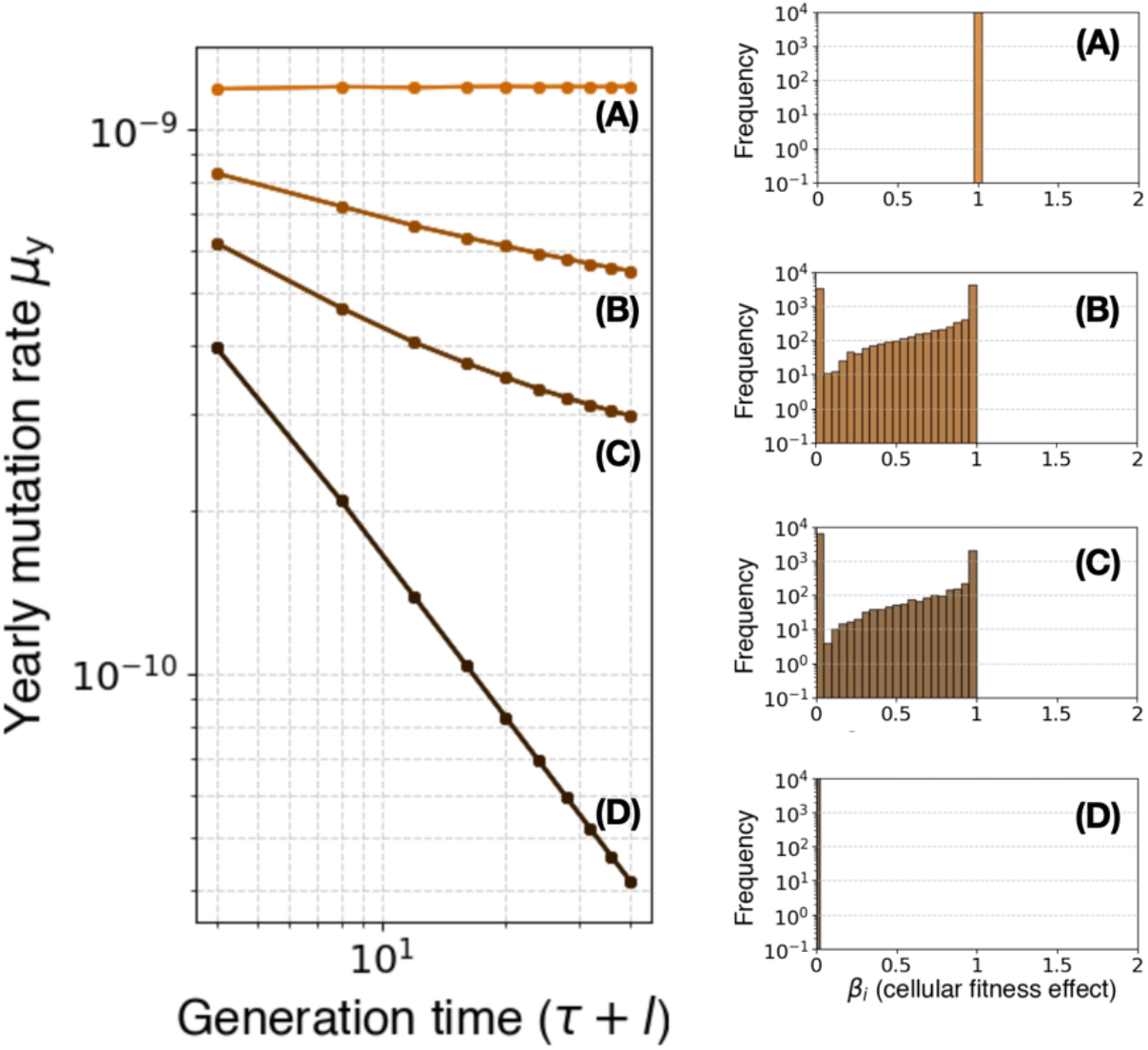
Cell-lineage selection may contribute to the anticorrelation between yearly mutation rates and generation time. Relationship between yearly mutation rates, *µ*_*y*_, and generation time in the logarithmic scale. *µ*_*y*_ was calculated as generational mutation rates, *µ*_*pop*_, divided by total generation time, consisting of the sum of developmental time, *τ*, and time to reproduction, *l*. Each curve represents the values calculated for the cellular distribution of fitness effects, cDFE, corresponding to each histogram A-D. The model parameters used for the results shown correspond to *β*_*i*_ = 1, *µ*_*mol*_ = 1*x*10^−10^, *C* = 1, *γ*=1, *d* = 0.05, and 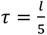, for different values of *l*. For each cDFE we considered our baseline shape parameters (Methods) with different proportions of deleterious and neutral mutations given by 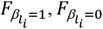, and 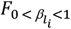. In (A) 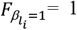, in (B) 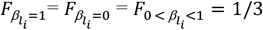, in (C) 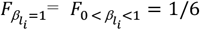 and 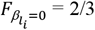, and in (D) 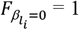. Note how for each of these scenarios we considered *l* to be directly proportional to *τ*. This is to isolate the effect of differences in proportions in developmental time, which may also contribute to the generation time effect (78).

**Figure S5.**
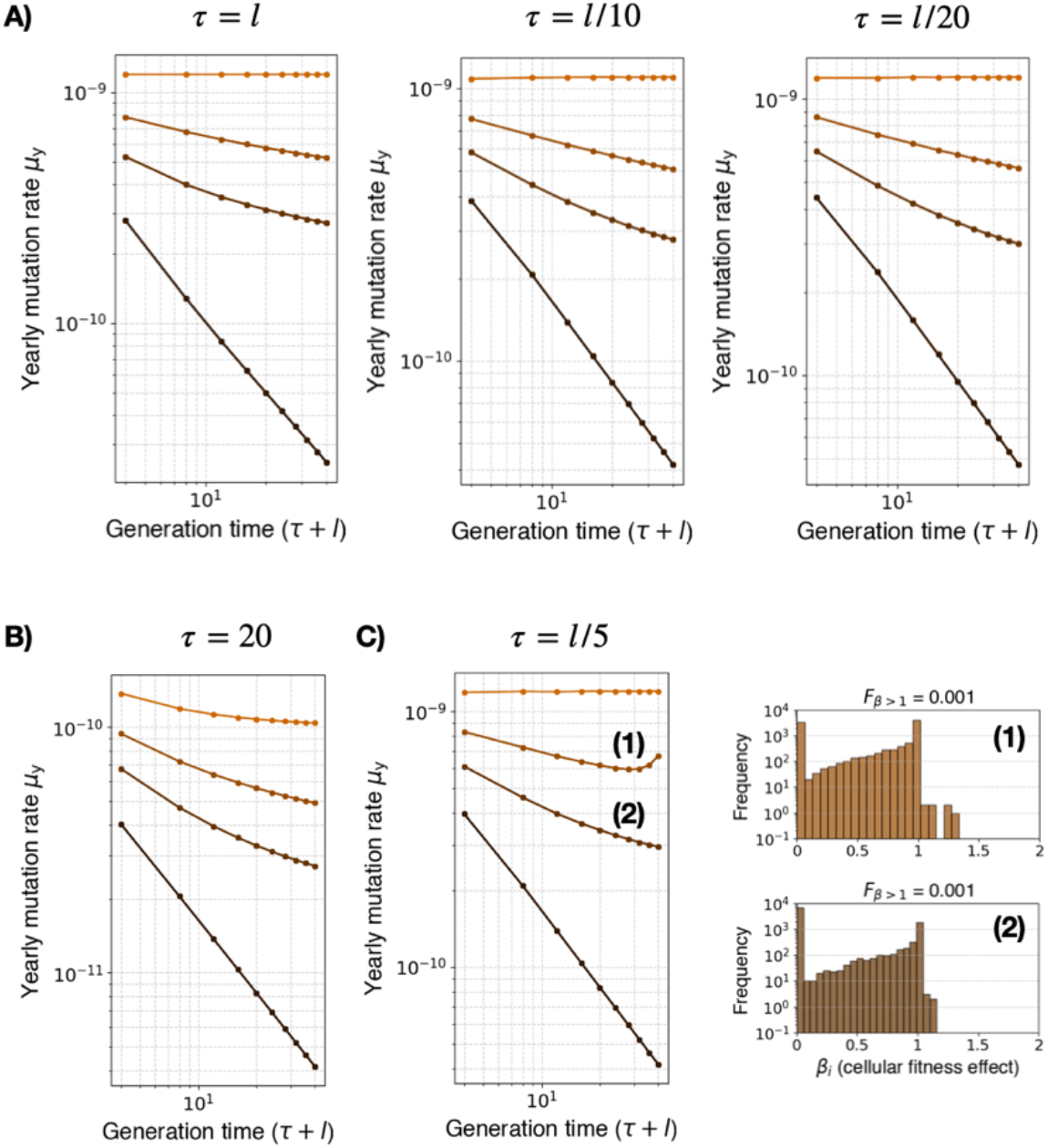
The anticorrelation between yearly mutation rates and generation time is robust to assumptions. (A) Relationship between yearly mutation rates, *µ*_*y*_, and generation time in the logarithmic scale. *µ*_*y*_ was calculated as generational mutation rates, *µ*_*pop*_, divided by total generation time, consisting of the sum of developmental time, *τ*, and time to reproduction, *l*. Each curve represents the values calculated for the cellular distribution of fitness effects, cDFE, corresponding to each histogram A-D shown in Figure S4. For each panel the dependence of *τ* on *l* was changed as indicated. The model parameters used for the results shown correspond to *β*_*i*_ = 1, *µ*_*mol*_ = 1*x*10^−10^, *C* = 1, *γ*=1, *d* = 0.05. For each cDFE we considered our baseline shape parameters (Methods) with different proportions of deleterious and neutral mutations given by 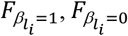, and 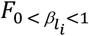. In (A) 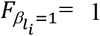, in (B) 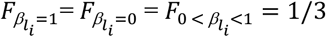, in 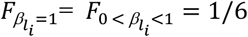 and 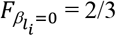, and in (D) 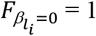. (B) Relationship between yearly mutation rates, *µ*_*y*_, and generation time in the logarithmic scale considering a non-proportionality between *τ* and *l. τ* was held constant with a value of 20. All other parameters are as indicated for (A). (C) Relationship between yearly mutation rates, *µ*_*y*_, and generation time in the logarithmic scale considering a proportionality as in Figure 5. In this case for the estimations indicated as (1) and (2) we considered also a fraction of mutations with 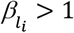, such that 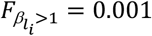. The respective cDFE for these cases are shown in the histograms.

